# Detection of a geminate photoproduct of bovine cytochrome c oxidase by time-resolved serial femtosecond crystallography

**DOI:** 10.1101/2023.05.08.539888

**Authors:** Izumi Ishigami, Sergio Carbajo, Nadia Zatsepin, Masahide Hikita, Chelsie E. Conrad, Garrett Nelson, Jesse Coe, Shibom Basu, Thomas Grant, Matthew H. Seaberg, Raymond G. Sierra, Mark S. Hunter, Petra Fromme, Raimund Fromme, Denis L. Rousseau, Syun-Ru Yeh

**Affiliations:** Department of Biochemistry, Albert Einstein College of Medicine, Bronx, NY, 10461, USA; Linac Coherent Light Source, SLAC National Accelerator Laboratory, Menlo Park. CA, 94025, USA; Electrical & Computer Engineering Department, University of California Los Angeles, Los Angeles, CA 90045; Physics & Astronomy Department, University of California Los Angeles, Los Angeles, CA 90045; Department of Physics, Arizona State University, Tempe, AZ 85287, USA; Center for Applied Structural Discovery, The Biodesign Institute, Arizona State University, Tempe, AZ, 85287, USA; Chemistry and Physics, La Trobe University, Kingsbury Drive, Bundoora, VIC, 3086, Australia; School of Molecular Sciences, Arizona State University, Tempe, AZ 85287, USA; Department of Structural Biology, University Buffalo, 955 Main Street, Buffalo, New York 14203, USA

## Abstract

Cytochrome *c* oxidase (C*c*O) is a large membrane-bound hemeprotein that catalyzes the reduction of dioxygen to water. Unlike classical dioxygen binding hemeproteins with a heme *b* group in their active sites, C*c*O has a unique binuclear center (BNC) comprised of a copper atom (Cu_B_) and a heme *a*_3_ iron, where O_2_ binds and is reduced to water. CO is a versatile O_2_ surrogate in ligand binding and escape reactions. Previous time-resolved spectroscopic studies of the CO complexes of bovine C*c*O (bC*c*O) revealed that photolyzing CO from the heme *a*_3_ iron leads to a metastable intermediate (Cu_B_-CO), where CO is bound to Cu_B_, before it escapes out of the BNC. Here, with a time-resolved serial femtosecond X-ray crystallography-based pump-probe method, we detected a geminate photoproduct of the bC*c*O-CO complex, where CO is dissociated from the heme *a*_3_ iron and moved to a temporary binding site midway between the Cu_B_ and the heme *a*_3_ iron, while the locations of the two metal centers and the conformation of the Helix-X, housing the proximal histidine ligand of the heme *a*_3_ iron, remain in the CO complex state. This new structure, combined with other reported structures of bC*c*O, allows the full definition of the ligand dissociation trajectory, as well as the associated protein dynamics.

## Main Text

Cytochrome c oxidase (C*c*O) is the terminal enzyme in the electron transfer chain in the inner membrane of mitochondria. It reduces dioxygen to two water molecules by accepting four electrons from cytochrome *c* and four protons from the negative side of the mitochondrial membrane. At the same time, it harnesses the chemical energy derived from the dioxygen reduction chemistry to translocate four protons from the negative to positive side of the membrane for the production of the electrochemical proton gradient required for ATP synthesis by ATP synthase.^1^ To accomplish this complex task, C*c*O possesses four redox active centers, Cu_A_, heme *a*, and a binuclear center (BNC) formed by a copper atom (Cu_B_) and the heme *a*_3_ iron,^2^ where O_2_ binds and is reduced to water. The dioxygen reduction reaction catalyzed by C*c*O has been extensively studied.^3-7^ It is thus relatively well-understood. In contrast, the O_2_ binding dynamics preceding the oxygen chemistry remains elusive as it is technically challenging to monitor the fleeting C*c*O-O_2_ complex.

Carbon monoxide (CO), like O_2_, is an excellent ligand for hemeproteins, but, unlike O_2_, it forms nonreactive complexes with them. Furthermore, when CO is bound to a heme iron, it is readily dissociable by a short pulse of visible light, making it a versatile tool for the investigation of ligand dissociation and rebinding dynamics.^8,9^ Time-resolved Fourier-transform infrared spectroscopic studies at ambient temperatures showed that photolyzing CO from the heme *a*_3_ iron in the BNC of bovine C*c*O (bC*c*O) in free solution leads to the formation of a characteristic Cu_B_-CO intermediate (Fig. 1A), where the photolyzed CO is coordinated to the Cu_B_, in less than 1 picosecond.^10-14^ The photolyzed CO subsequently disociates from Cu_B_ and escapes out of the BNC in ∼1.5 μs,^11,15^ to generate the ligand-free reduced species (R). By extention, it is believed that the bimolecular binding reaction of CO, as well as O_2_, follows the same reaction trajectery, but in a reverse order, with Cu_B_ as a temporary ligand stop.^6,16-18^ In addition to the Cu_B_-CO intermediate, transient UV-Vis absorption spectroscopic data suggest that a geminate photoproduct, where CO has been photolyzed from heme *a*_3_ iron but not yet coordinated to Cu_B_, is formed immediately following CO photodissociation.^11,15^ This proposed intermediate, however, has never been experimentally identified.

**Fig. 1.**
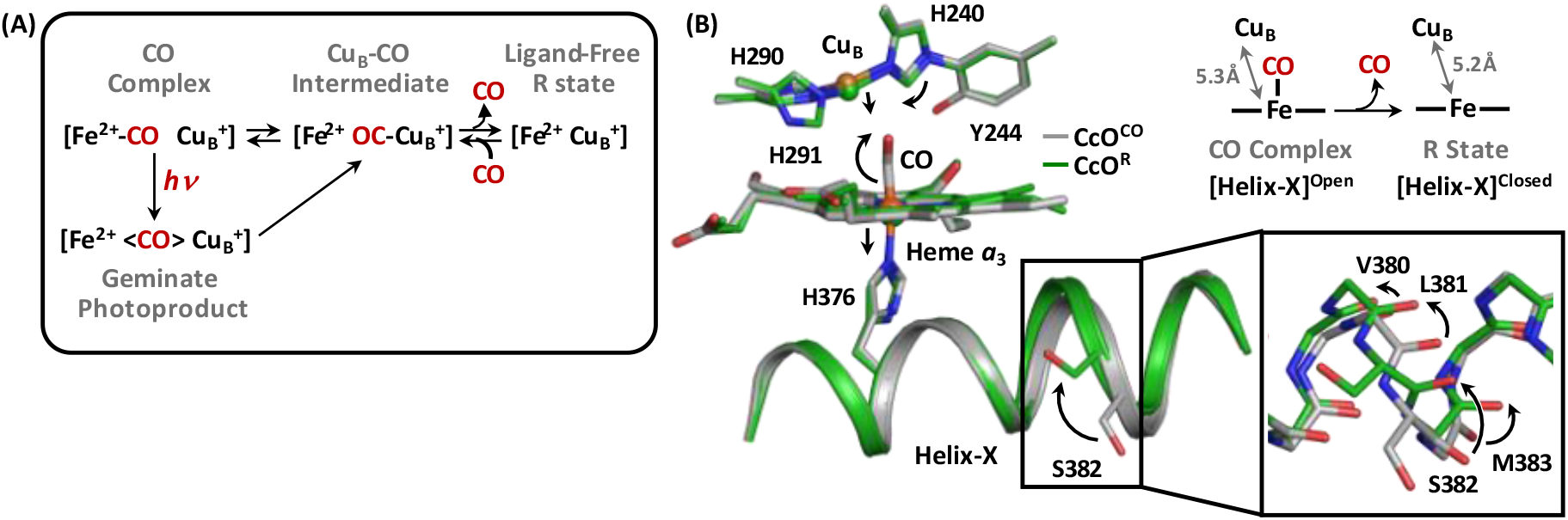
Proposed CO dissociation mechanism of bC*c*O (A) and the associated structural transition (B). The structure of the CO complex (PDB ID: 5W97) (grey) is superimposed with that of the ligand-free R species (PDB ID: 7THU) (green) in (B) to illustrate the CO dissociation induced structural transition (as indicated by the arrows), in particular (i) the out-of-plane movement of the heme *a*3 iron-H376 moiety, (ii) the displacement of the CuB-H240 moiety towards the heme *a*3, and (iii) the open to closed conformational change in Helix-X. The inset shows the expanded view of the [380-383] fragment of Helix-X. The arrows indicate the rotation of the backbone carbonyl groups. The sidechains, except that of S382, are not shown for clarity.

Crystallographic studies of the bC*c*O-CO complex showed that X-rays from synchrotron light sources, like visible light, can photolyze CO from heme *a*_3_.^19^ To prevent the X-ray induced radiation damage, we employed serial femtosecond crystallography (SFX)^20^ to determine the structure of the intact bC*c*O-CO complex.^19^ With SFX, the diffraction patterns of randomly oriented microcrystals suspended in an aqueous jet, which intersects the femtosecond pulses from a X-ray free-electron laser (XFEL), are sequentially collected and then merged for structural determination. As the raditaion damage processes do not occur until after the termination of each ultrashort laser pulse, radiation damage-free structures can be obtained under native-like conditions at ambient temperatures.^20^ The comparison of the SFX structure of the bC*c*O-CO complex with that of the ligand-free protein (R) (Fig. 1B) reveals that CO dissociation leads to the displacement of the Cu_B_-H240 moiety towards the heme *a*_3_ and the movement of the heme *a*_3_ iron, as well as the proximal histidine ligand (H376) coordinated to it, away from the heme *a*_3_ plane. The structural changes to the BNC are associated with the conversion of the Helix-X housing H376 from an open to a closed conformation,^21^ where the [380-383] backbone rotates from a bulged structure to an α-helical structure and the S382 sidechain flips by ∼180°.

To comprehend the protein dynamics associated with CO dissociation reaction of the bC*c*O-CO complex, here we sought to employ time-resolved SFX-based pump-probe method to determine the structure of an early photoproduct of the bC*c*O-CO complex, using a tunable-wavelength optical parametric oscillator (OPO) as the pump beam to photodissociate CO and a XFEL as the probe beam to monitor the structural changes. The suspension of the plate-like microcrystals (∼20x20x4 μm in size), prepared based on a previously reported protocol,^19^ was loaded into a gas-tight syringe and injected into the XFEL beam with a gas dynamic virtual nozzle (GDVN) injector^22^ as a thin (4 μm) aqueous solution jet in the vacuum chamber at the CXI experimental station at the Linac Coherent Light Source (LCLS) at the SLAC National Accelerator Laboratory. The output wavelength and pulse energy from the OPO laser were set at 492 nm and 60 mJ, respectively, to ensure that the CO ligand was completely photolyzed. Each OPO laser pulse (with a pulse width of approximately 8 ns) was timed at 100 ns before the XFEL pulse (with a pulse width of ≤ 40 fs) by a precise timing synchronization system. The serial diffraction patterns were collected for 85 mins, among which 16,520 indexable patterns were merged and analyzed.

The initial structure was solved with molecular replacement using the structure of ligand-free reduced bC*c*O (PDB ID: 7THU) as the search model. In the F_O_-F_C_ difference map (contoured at 6 σ) (see inset-i in Fig. 2A), clear ligand electron density is evident between Cu_B_ and heme *a*_3_; in addition, there is no residual electron density connecting the ligand density to the heme *a*_3_ iron (even when the map is contoured at 3σ), indicating that CO is completely photolyzed and that there is no geminate CO recombination, in good agreement with free solution reactions.^23^ As the C and O atoms are indistinguishable in the current data, the ligand electron density was modeled with a CO molecule with the atom closer to the iron arbitrarily assigned as the C atom (inset-ii). The final structure was refined to a resolution of 2.8 Å (PDB ID: 8GBT, see Extended Data Table 1).

**Fig. 2.**
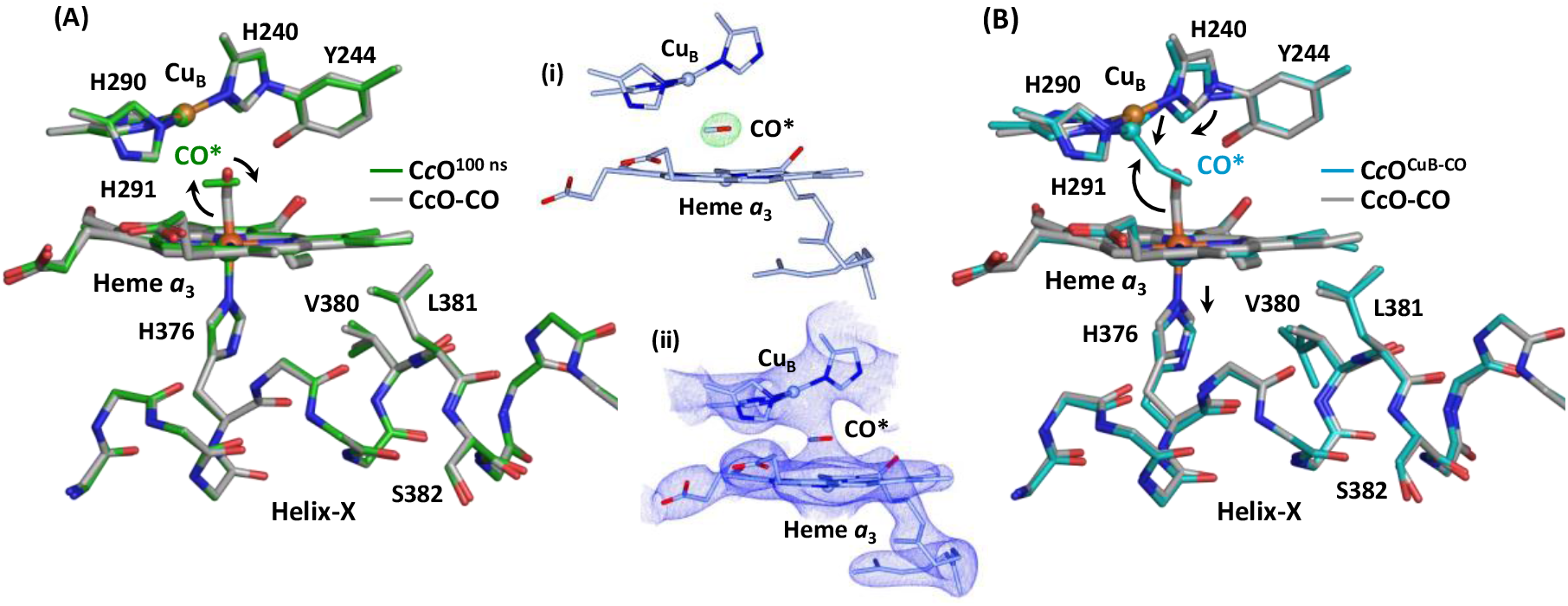
Structure of the geminate photoproduct derived from this work (PDB ID: 8GBT) (A) and that of the reported CuB-CO intermediate (PDB ID: 5X1B) (B). The structures are superimposed with that of the bC*c*O-CO complex (PDB ID 5W97) (grey) to highlight the structural differences between the two photoproducts. In the geminate photoproduct reported here, the CO is photolyzed and moved to a new position midway between heme *a*3 and CuB, while the BNC and the Helix-X remain in the CO-complex state; in contrast, in the CuB-CO intermediate, CO coordinates to CuB, the heme *a*3 iron-H376 moiety moves out of the heme *a*3 plane and the CuB-H240 moiety displaces towards the heme *a*3 (as indicated by the arrows), while the Helix-X remains in the CO-complex (open) state. The inset (i) in (A) displays the FO-FC difference map of the BNC (contoured at 6 σ), showing the clear electron density associated with CO between heme *a*3 and CuB in the BNC. The inset (ii) shows the 2FO-FC map of the BNC (contoured at 2 σ) with the ligand electron density modeled with CO.

The structural data reveal that, upon photodissociation, the CO rotates ∼90° with respect to its center of mass, such that it lies parallel with the heme *a*_3_ plane, midway between Cu_B_ and the heme *a*_3_ iron. The Cu_B_-H240 and heme *a*_3_ iron-H376 moieties, as well as the conformation of the Helix-X, remain unperturbed, indicating that the proten matrix does not relax to the ligand-free R state until later during the reaction. The structure is distinct from that of a photoproduct reported by Shimada *et al* (PDB ID: 5X1B),^24^ derived from a different pump-probe experiment, where CO is photolyzed from the heme *a*_3_ iron and coordinated to Cu_B_ (Fig. 2B). In that state, the Cu_B_-H240 moiety moved towards the heme *a*_3_ iron by 0.6 Å and the heme *a*_3_ iron-H376 moiety moved away from the heme plane by 0.2 Å, resulting in the contraction of the Cu_B_-heme *a*_3_ iron distance from 5.3 to 5.0 Å, while the Helix-X remained in the CO-complex (open) state. The structural features of the two photoproducts, combined with the reaction scheme illustrated in Fig. 1A, suggest that the 100 ns intermediate reported in this work is the long-suspected primary geminate photoproduct, while the Shimada structure is the Cu_B_-CO intermediate. It is important to note that the two photoproducts are distinct from that generated during X-ray diffraction data collection induced by constant synchrotron light illumination (PDB ID: 5WAU)^19^ (see Fig. S1 in the Supplemental Data), as the latter represents a unique photostationary state, instead of a true reaction intermediate.

The availability of the structures of the two reaction intermediates of bC*c*O, combined with the equilibrium structures of the CO-complex and ligand-free R state, allows us to define the ligand dissociation trajectory and the associated protein dynamics in bC*c*O as depicted in Fig. 3. The heme *a*_3_ iron-CO bond scission first leads to the rotation of the CO to a new orientation parallel with the heme *a*_3_ plane (step 1, indicated by the green arrow in Fig. 3B), while the Cu_B_-H240 and heme *a*_3_ iron-H376 moieties and the Helix-X remain in the CO-complex (open) state. In the ensuing reaction (step 2), CO further rotates towards Cu_B_ to establish a coordination bond with it. It triggers the relaxation of the BNC to a ligand-free R-like conformation (see the blue arrows), cumulating in 0.3 Å contraction of the Cu_B_-heme *a*_3_ iron distance, while the conformation of the Helix-X remains unperturbed. Finally, CO dissociates from Cu_B_ and escapes out of the BNC, accompanied by a 0.2 Å re-expansion of the Cu_B_-heme *a*_3_ iron distance and the subsequent rotation of the [380-383] peptide backbone of the Helix-X to the equilibrium R (closed) conformation (step 3, indicated by the red arrows), thereby completing the reaction.

**Fig. 3.**
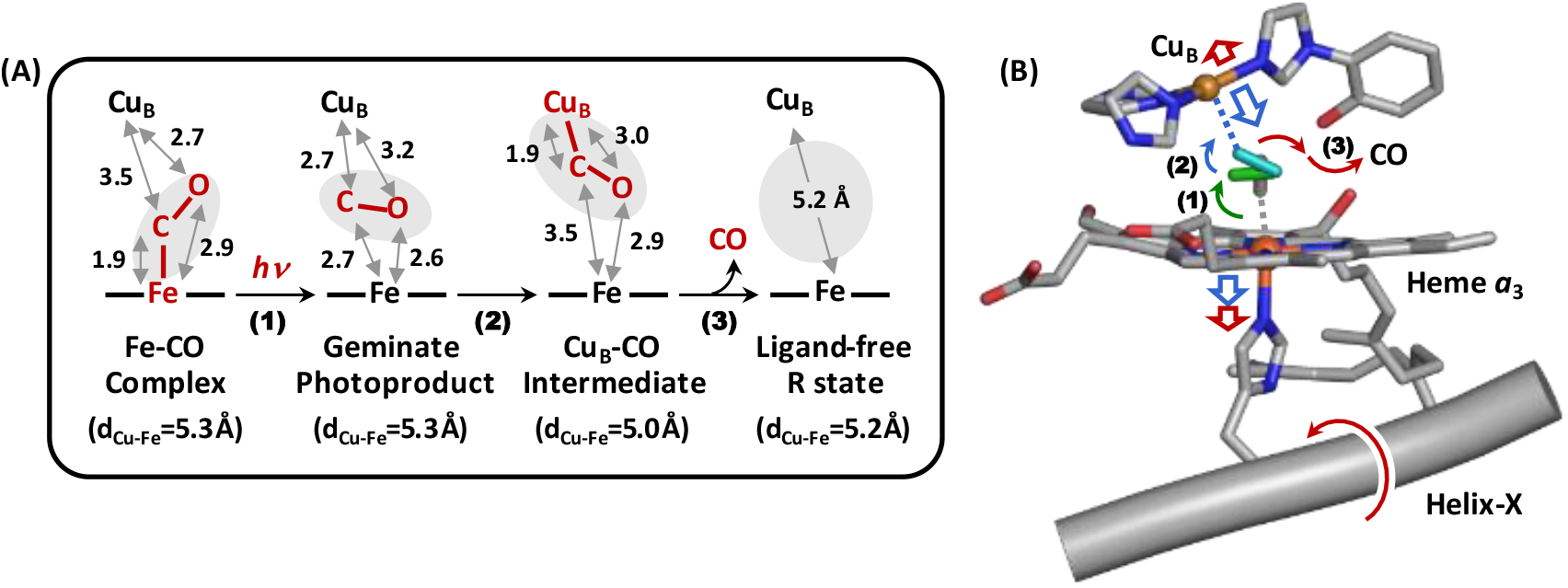
CO dissociation reaction trajectory in bC*c*O. (A) The three elementary steps of the CO dissociation reaction and the interatomic distances in each species involved in the reaction. (B) The bC*c*O-CO complex structure (PDB ID 5W97) (grey) superimposed with the CO molecules taken from the geminate photoproduct (this work) (green) and the CuB-CO complex (PDB ID: 5X1B) (cyan). The green, blue and red arrows indicate the conformational transition associated with the step (1), (2) and (3) of the reaction, respectively.

The overall protein structural transition induced by CO dissociation in bC*c*O is comparable to that in myoglobin (Mb), a model system considered as the hydrogen atom of biology.^25^ Mb is a small water soluble protein that contains a heme *b* prosthetic group embedded in the protein matrix. The heme iron is coordinated by a histidine residue (H93) from Helix-F on the proximal side (equivalent to Helix-X in bC*c*O), and a diatomic ligand, such as O_2_ or CO, on the distal side, which is stabilized by another histidine residue (H64) from the Helix-E (in place of the Cu_B_ moiety in bC*c*O). Photodissociation of CO from the Mb-CO complex leads to a geminate photoproduct, where CO rotates and translates to a nearby docking site, such that it lies parallel with the heme plane as in bC*c*O, within the distal heme pocket.^26^ It is accompanied by the displacement of the heme iron out of the heme plane, as well as the concurrent movement of (i) the proximal H93 and the associated Helix-F away from the heme, and (ii) the distal H64 and the associated Helix-E towards the heme.^27-29^. These structural changes near the active site then propagate to the rest of the protein matrix through a global protein relaxation in a quake-like motion to establish the final equilibrium ligand-free structure.^27-29^

Despite the similarities, the protein dynamics associated with CO dissociation in bC*c*O is much less cooperative as compared to that in Mb. The dissociation of CO from the heme iron in bC*c*O, like that in Mb, leads to the formation of a geminate photoproduct, with CO bound to a docking site within the distal heme pocket, but it does not introduce any immediate structural changes to the heme or the protein matrix surrounding it. The presence of Cu_B_ in the BNC in bC*c*O reroutes the CO migration pathway, rendering an additional Cu_B_-CO intermediate. The migration of CO from the docking site to Cu_B_ triggers the displacement of the distal Cu_B_-H240 moiety and the out-of-plane movement of the heme *a*_3_ iron-H376 moiety, while the subsequent escape of CO out of the BNC induces the final relaxation of the proximal Helix-X to the final equilibrium R state.

These unique CO dissociation-induced protein dynamics in bC*c*O is accompanied by transient contraction of the BNC (see Fig. 3A), which plausibly plays an important role in guiding the unidirectional movement of the ligand out of the BNC, thereby accounting for the lack of geminate recombination in the bC*c*O reaction, in contrast to the ∼40% geminate recombination in the Mb reaction.^27^ This characteristic protein plasticity in bC*c*O is conceivably critical for coupling the oxygen chemistry occurring in the BNC to proton translocation.

In summary, the data presented here provide the first direct experimental structural evidence of the presence of the primary geminate photoproduct during the CO photodissociation reaction of bC*c*O. The definition of its structure, combined with a previously reported Cu_B_-CO structure (PDB ID: 5X1B)^24^, enables us to define the entire ligand dissociation trajectory, as well as the associated protein dynamics, for the first time, thereby offering exciting experimental blueprints for computational interrogation of the energy landscape associated with the ligand migration reaction.

## Acknowledgements

The SFX experiments were carried out at the Linac Coherent Light Source (LCLS) at the SLAC National Accelerator Laboratory. LCLS is an Office of Science User Facility operated for the US Department of Energy Office of Science by Stanford University. Use of the LCLS, SLAC National Accelerator Laboratory, is supported by the US Department of Energy, Office of Science, Office of Basic Energy Sciences, under Contract DE-AC02-76SF00515. Parts of the sample delivery system used at LCLS for this research were funded by NIH Grant P41GM103393, formerly P41RR001209. This work was supported by National Science Foundation (NSF) grants CHE-1404929 (to D.L.R. and S.-R. Yeh), ABI Innovations Award 1565180 (to N.A.Z.) and Science and Technology Center Award 1231306. This work was also supported by National Institutes of Health (NIH) grants GM126297 (D.L.R. and S.-R.Y.), GM115773 (S.-R.Y.), GM095583 (to P.F.), S10 OD023453 and P41 GM139687, as well as the Biodesign Center for Applied Structural Discovery at Arizona State University.

## Author contributions

I.I., S.C., S.-R.Y., and D.L.R. designed research; I.I. isolated and crystallized bCcO; I.I., M.H., C.E.C.,G.N., J.D.C., M.H.S., R.G.S., M.S.H., and R.F. performed research; S.-R.Y. contributed new analytic tools; N.A.Z., S.B., T.D.G., R.F., S.-R.Y., and D.L.R. analyzed data; and S.-R.Y. and D.L.R. wrote the initial draft of the manuscript.

