## Supporting Information for "Detection of a geminate photoproduct of bovine cytochrome c oxidase by time-resolved serial femtosecond crystallography"

---

[1] Present address: Masahide Hikita, Structural Biology Research Center, Institute of Materials Structure Science, High Energy Accelerator Research Organization, 1-1 Oho, Tsukuba, Ibaraki 305-0801 Japan.
[2] Present address: Chelsie E. Conrad, Center for Genomic Medicine, University of Utah, 30 South 2000 East, Salt Lake City, Utah 84112, USA.
[3] Present address: Garrett Nelson, Boeckeler Instruments, Inc. 4650 S Butterfield Dr, Tucson, AZ 85714, USA.
[4] Present address: Jesse Coe, KBI Biopharma, 1101 Hamlin Rd, Durham, NC 27704, USA.
[5] Present address: Shibom Basu, EMBL, Grenoble, 71 Avenue des Martyrs, CS 90181, 38042 Grenoble, France.

### Materials and Methods

*Microcrystal preparation.* Bovine C*c*O was isolated from bovine hearts by a standard procedure.[1,2] In the final step of the purification, the enzyme was crystalized by concentration on an Amicon concentrator equipped with a 200 kDa molecular cutoff filter. The crude crystals were harvested and redissolved in 40 mM pH 6.8 phosphate buffer containing 0.2% decylmaltoside to generate the protein stock. The microcrystals were prepared with a previously reported method.[3] The crystal growth was initiated by mixing the protein stock with the precipitant solution (0.2% decylmaltoside and 2.5% PEG4000 in 40 mM pH 6.8 phosphate buffer) and a seeding solution (prepared by crushing and sonicating large crystals in the mother solution). The microcrystals were allowed to grow at 4 $^{o}$C for ~36 hours before they were harvested and characterized by polarized optical microscopy. The microcrystals have a planar shape with approximate dimensions of ~20 x 20 x 4 μm. To generate the CO-complex, the microcrystals were purged with $N_2$, reduced with minimum amount of dithionite, and then exposed to CO.

*Time-resolved SFX data collection and analysis.* As reported previously, the tr-SFX experiments were carried out at the CXI experimental endstation at the Linac Coherent Light Source (LCLS) at the SLAC National Accelerator Laboratory.[3] The suspension of the bC*c*O microcrystals was loaded into a gas-tight syringe and injected out of a gas dynamic virtual nozzle (GDVN) injector as a thin solution jet at a flow rate of ~10 µl/min in the vacuum chamber at the CXI station. A 492 nm beam from an OPO (60 uJ, 8 ns pulse width, 120 Hz maximum repetition rate), with a FWHM transverse beam size of 250 μm, was used as the pump beam to photolyze CO, and the 9.5 keV XFEL beam (≤40 fs pulse width, 120 Hz), with a FWHM transverse beam size of ~1 μm, was used as the probe beam to obtain the diffraction patterns. The two beams were aligned perpendicular to the solution jet and timed using a precise timing synchronization system with $\Delta t = 100$ ns.

The diffraction patterns were collected by a Cornell-SLAC Pixel Array Detector (CSPAD) at a 120 Hz rep rate. The quality and hit rate of the SFX data were monitored in real time using OnDA[4]. Cheetah[5] was used for detector corrections and hit finding (data reduction). *CrystFEL*'s indexing program was used to index the crystal hits and merge the patterns.[6,7] A total of 16,520 indexable diffraction patters were obtained. The initial structure was solved with molecular replacement with Phaser-MR through the CCP4 program suite[8] using the structure of ligand-free reduced bC*c*O (PDB ID: 7THU) as the search model. Further model building was performed using Coot.[9] Structure refinements were done using Refmac5 and PDB-Redo[10]. The final structure was refined to a resolution of 2.38 Å (see Extended Data Table 1) (PDB ID: 8GBT).

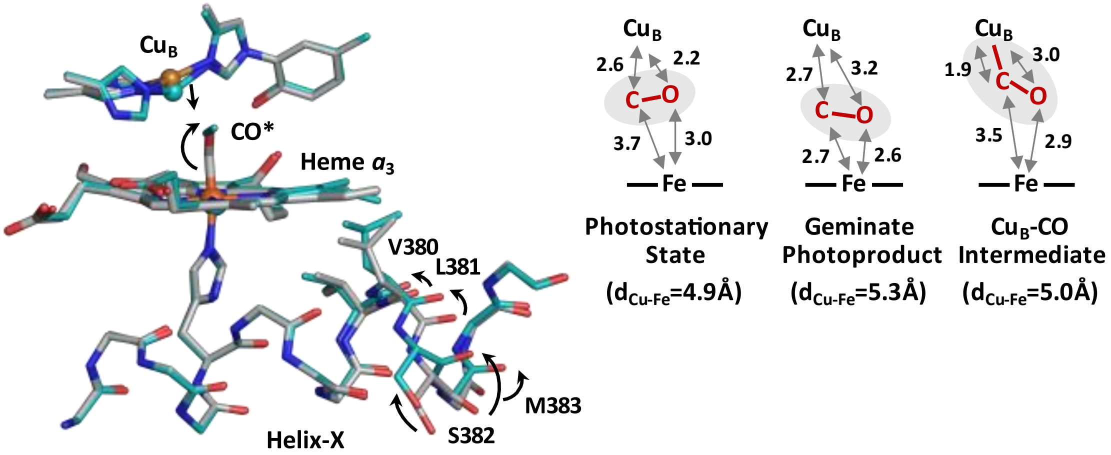


**Fig. S1. Structure of a photoproduct of the bC*c*O-CO complex induced by synchrotron light illumination (PDB ID: 5WAU)[3].** The structure of the photoproduct (cyan) is overlaid with that of the intact bCcO-CO complex (grey) to illustrate the photolysis of CO and the associated conformational changes, in particular (i) the 0.5 Å downward displacement of the $Cu_B$ towards the heme $a_3$ (hence a contraction of the $Cu_B$-heme $a_3$ iron distance from 5.3 to 4.9 Å) and (ii) the conversion of the [380-383] peptide backbone in Helix-X to a unique ligand-free R like (closed) conformation. The structure of this photoproduct is distinct from that of the geminate photoproduct or the $Cu_B$-CO intermediate (see Fig. 2 in the main text), indicating that the constant synchrotron X-ray illumination during the diffraction data acquisition converts the protein to a unique photostationary state. The right inset shows the comparison of the interatomic distances in the BNC of the photostationary state with those in the BNC of the geminate photoproduct reported here and the $Cu_B$-CO intermediate reported by Shimada *et al*.[11] It's notable that both the geminate photoproduct and the $Cu_B$-CO intermediate were generated by time-resolved methods, but the difference in the photolysis wavelength (492 nm *versus* 532 nm) and intensity (60 µJ in a 8 ns pulse *versus* 200 µJ in a 4 ns pulse), as well as the crystal size (microcrystals *versus* large single crystals), plausibly leads to the difference in the CO migration kinetics.

**Extended Data Table 1**. Crystallographic data collection and refinement statistics (PDB ID: 8GBT). The values in the parentheses are for the outer shell.

| **Data Collection** | |
|---|---|
| Space Group | $P2_12_12_1$ |
| Dimensions: a, b, c (Å) | 178.3, 189, 209 |
| Dimensions α, β, γ (°) | 90, 90, 90 |
| Resolution (Å) | 32.00 – 2.8 |
| I/σI | 6.92 (0.43) |
| CC* | 0.9612 (0.607) |
| Redundancy | 370.3 |
| Completeness (%) | 100 (100) |
| Number of Indexed Hits | 16,520 |
| **Refinement** | |
| Resolution (Å) | 32.00 – 2.8 |
| Unique reflections | 172,414 (17,697) |
| $R_{work}/R_{free}$ | 0.223/0.2687 (0.36/0.372) |
| Number of atoms | 30,981 |
| Average B factor (Å$^2$) | 58.179 |
| **R.m.s. deviations** | |
| Bond lengths (Å) | 0.006 |
| Bond angles (°) | 1.553 |
| **Ramachandran statistics (%)** | |
| Favored regions | 89.84 |
| Outliers | 1.35 |
| Molprobity score | 2.48 |
